# B1L regulates lateral root development by exocytic vesicular trafficking-mediated polar auxin transport in Arabidopsis

**DOI:** 10.1101/2021.06.07.447418

**Authors:** Gang Yang, Bi-xia Chen, Tao Chen, Jia-hui Chen, Rui Sun, Cong-cong Liu, Jiao Jia, Xiu-le Yue, Li-zhe An, Hua Zhang

## Abstract

Auxin and auxin-mediated signaling pathways involved in the regulation of lateral root development are well documented. Although exocytic vesicle trafficking plays an important role in auxin efflux carriers PIN recycling, and polar auxin transport during lateral root formation, however, the mechanistic details of these processes are not well understood. Here, we demonstrate that BYPASS1-LIKE (B1L) regulate lateral root development via exocytic vesicular trafficking-mediated polar auxin transport in Arabidopsis. In *b1l* mutants, the number of lateral roots increased significantly, and the phenotypes were mainly attributed to lateral root primordium initiation but not to the defects in lateral root primordium development. Furthermore, the auxin signal was stronger in the lateral root primordium of the *b1l* mutant at stage I than those observed in the wild-type (WT). Moreover, exogenous auxin and auxin transport inhibitory treatments indicated that the phenotype of lateral roots in *b1l* mutants can be attributed to higher auxin levels and that B1L regulates auxin efflux. Consistently, auxin efflux carriers PIN1-GFP and PIN3-GFP were expressed at higher levels in the lateral root primordium of the *b1l* mutants. Interestingly, we found that B1L interacted with the exocyst and *b1l* mutant showed a defect in PIN2 exocytosis. Finally, we found that B1L cooperated with EXO70B1 to regulate lateral root formation. Our findings reveal an essential regulatory mechanism of B1L that interacts with the exocyst to regulate PIN-mediated polar auxin transport and lateral root initiation.

## INTRODUCTION

Roots are important vegetative organs that make plants sessile in soil and function in water and nutrient absorption (Bellini *et al*., 2014; Motte *et al*., 2019). Lateral root formation is a crucial event in root system morphogenesis(Lynch, 1995). This process is divided into five main steps: pre-branch site priming, lateral root initiation, lateral root patterning, lateral root emergence, and lateral root elongation(Banda *et al*., 2019; Péret *et al*., 2009). In turn, the phase between lateral root initiation and its patterning includes steps of the onset and formation of lateral root primordia. Generally, this phase is divided into eight stages (I-VIII) according to the layer number of lateral root primordial cells.(Stoeckle *et al*., 2018; Vilches-Barro and Maizel, 2015). Auxin and auxin-mediated signaling pathways play a central role in regulating all the steps of lateral root formation (Lavenus *et al*., 2013). Specifically, a local auxin threshold maintained by auxin influx carrier AUX1 (AUXIN RESISTANT1) and auxin efflux carriers PIN (PIN-FORMED) is essential for lateral root priming and initiation(Benková *et al*., 2003; Lavenus *et al*., 2013).

Furthermore, vesicle trafficking, which includes endocytosis and exocytosis, is an important physiological process widely involved in cell polarity establishment, cell expansion and division, cell wall formation, hormone signaling, and defense against pathogens(Wu *et al*., 2014; Zhang *et al*., 2019; Robatzek, 2007; Tanaka *et al*., 2006). Exocytic vesicle trafficking mainly includes targeting, tethering, and fusion(Zhang *et al*., 2019; Uemura and Ueda, 2014). An exocyst is a vital tethering factor that guarantees the specificity of contact between exocytic vesicles and the plasma membrane (Žárský *et al*., 2013; Mei and Guo, 2018; Wu and Guo, 2015). To date, eight subunits (SEC3, SEC5, SEC6, SEC8, SEC10, SEC15, EXO70, and EXO84) have been found in Arabidopsis.(Eckardt, 2008; Elias *et al*., 2003; Fendrych *et al*., 2010). Recycling of PIN1 and PIN2 is delayed in *exo70a1, sec8*, and *sec6* mutants(Drdová *et al*., 2013; Tan *et al*., 2016). These findings demonstrate that the exocyst plays an important role in PINs recycling and polar auxin transport. However, little is known about the precise mechanisms underlying these processes.

BYPASS1-LIKE (B1L) belongs to the DUF793 family whose functions are largely unknown. Our recent studies revealed that B1L interacts with 14-3-3λ to inhibit the degradation of CBF3 (C-REPEAT BINDING FACTOR3) and positively regulates freezing tolerance in Arabidopsis.(Chen *et al*., 2019). In addition, B1L interacts with TRANSTHYRETIN-LIKE (TTL) to regulate growth and freezing tolerance (Chen *et al*., 2020). However, to the best of our knowledge, no study has been yet reported on the other molecular functions of B1L. Here, we report a novel role of B1L revealed by using multiple approaches including, genetics, cellular biology, proteomics, and biochemical assays, namely, its involvement in lateral root development. B1L was revealed to be involved in exocytic vesicle trafficking-mediated PIN recycling through interaction with the exocyst and to regulate polar auxin transport to promote lateral root initiation. This study improves our understanding of the highly sophisticated processes involved in exocytic vesicular trafficking-mediated polar auxin transport and lateral root initiation in plants.

## RESULTS

### B1L is highly expressed in Arabidopsis roots

To gain further insight into the molecular function of B1L, we first examined its expression profile in roots, leaves, and whole seedlings. Western blot analysis revealed that the expression levels of B1L-3×Flag were significantly higher in roots than in leaves or whole seedlings (Fig. 1a, b). Next, we constructed a *B1L::GUS* transgenic plant to further investigate the expression pattern of B1L. GUS staining revealed that *B1L* was expressed extensively in various tissues; thus, particularly high levels of expression were observed in the vascular cylinder, in the meristematic zone of the primary root tissue, and at eight different developmental stages of lateral root primordia (Fig. 1c-m). These results indicated that B1L may be involved in root development.

**Figure 1.**
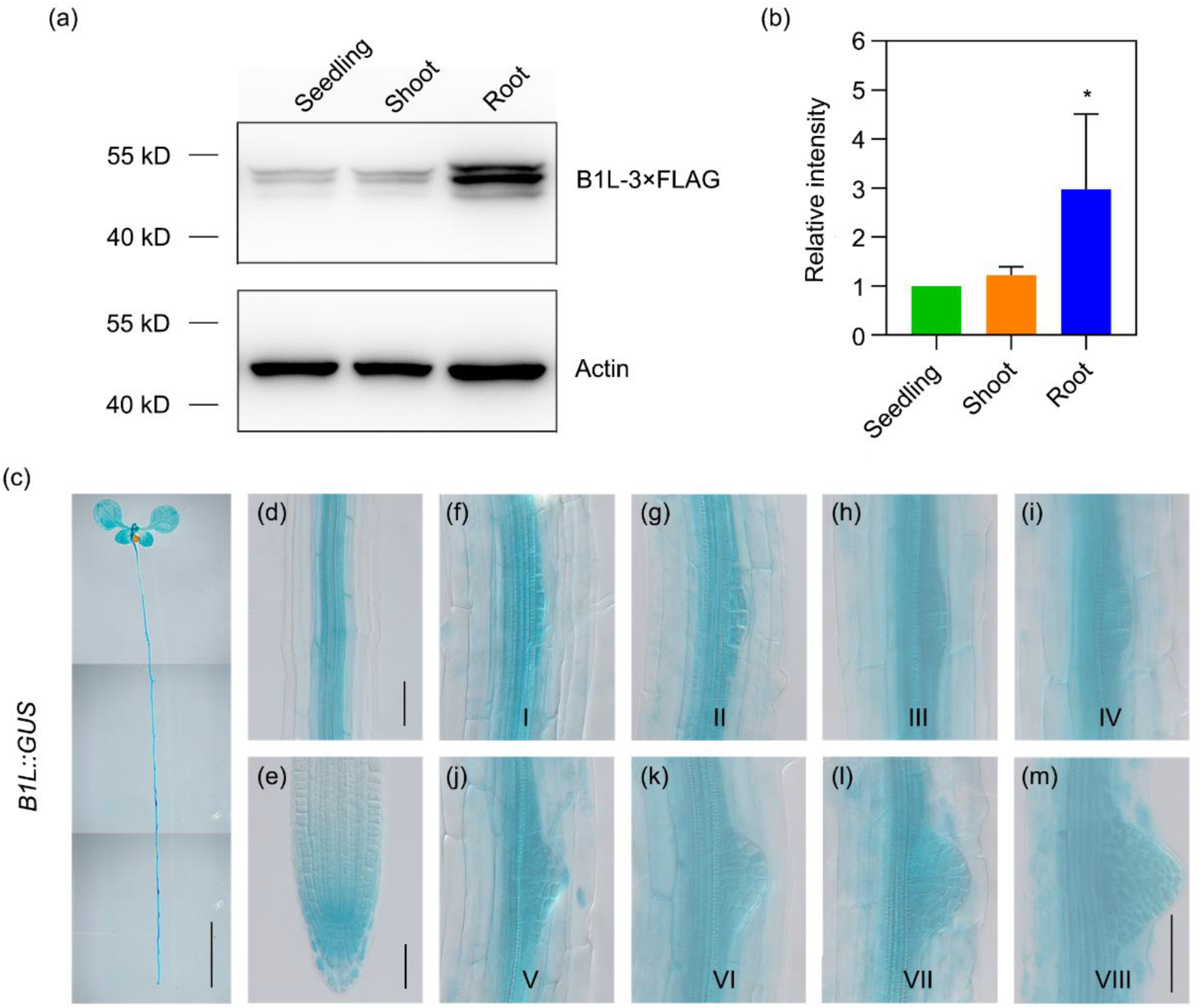
The expression pattern of B1L in *Arabidopsis* seedlings. (a) Western blot analysis of B1L in seeding, leaf and root of 10 days old *B1L::B1L-3×Flag/b1l* seedlings. B1L-3×Flag was recognized by the anti-Flag and Actin was used as an internal control. (b) Quantitative analysis of the B1L-3×Flag expression shown in A, with Image J software. Intensity of leaf was set as 1, the values are means ± SD (n=3, *one-way ANOVA*, *, P < 0.05). (c-m) GUS staining analysis of *B1l* expression in whole plant, primary root and eight different stages of lateral root primordium at 7 day after germination. I-VIII are eight development stages of lateral root primordia. Images were obtained by differential interference contrast microscope (DICM), bar in (c) is 5mm, and bars in (d-m) are 50μm.

### B1L regulates lateral root initiation

To examine whether B1L is involved in root development, WT, *b1l-1, b1l-2*, and *b1l-4* were grown on 1/2 MS medium plates under normal conditions and their phenotypes were monitored. We observed no significant difference in the length of primary roots among WT, *b1l-1, b1l-2*, and *b1l-4* (Fig. S2). However, the number of lateral roots in *b1l-1, b1l-2*, and *b1l-4* was significantly higher than that in the WT (Fig. 2a, b). Moreover, two independent transgenic lines expressing B1L-3×Flag driven by the native *B1L* promoter in *b1l-1*(*B1L::B1L-3×Flag/b1l*) exhibited the same number of lateral roots as the WT (Fig. 2 a, b), which indicated that the increase in the number of lateral roots in mutants was caused by the absence of B1L. In addition, two independent transgenic lines overexpressing *B1L* driven by the 35S promoter in the WT (*B1L*-OX#1 and *B1L*-OX#2) exhibited significantly fewer lateral roots than the WT (Fig. 2 a, b). These results demonstrated that B1L regulates lateral root formation.

**Figure 2.**
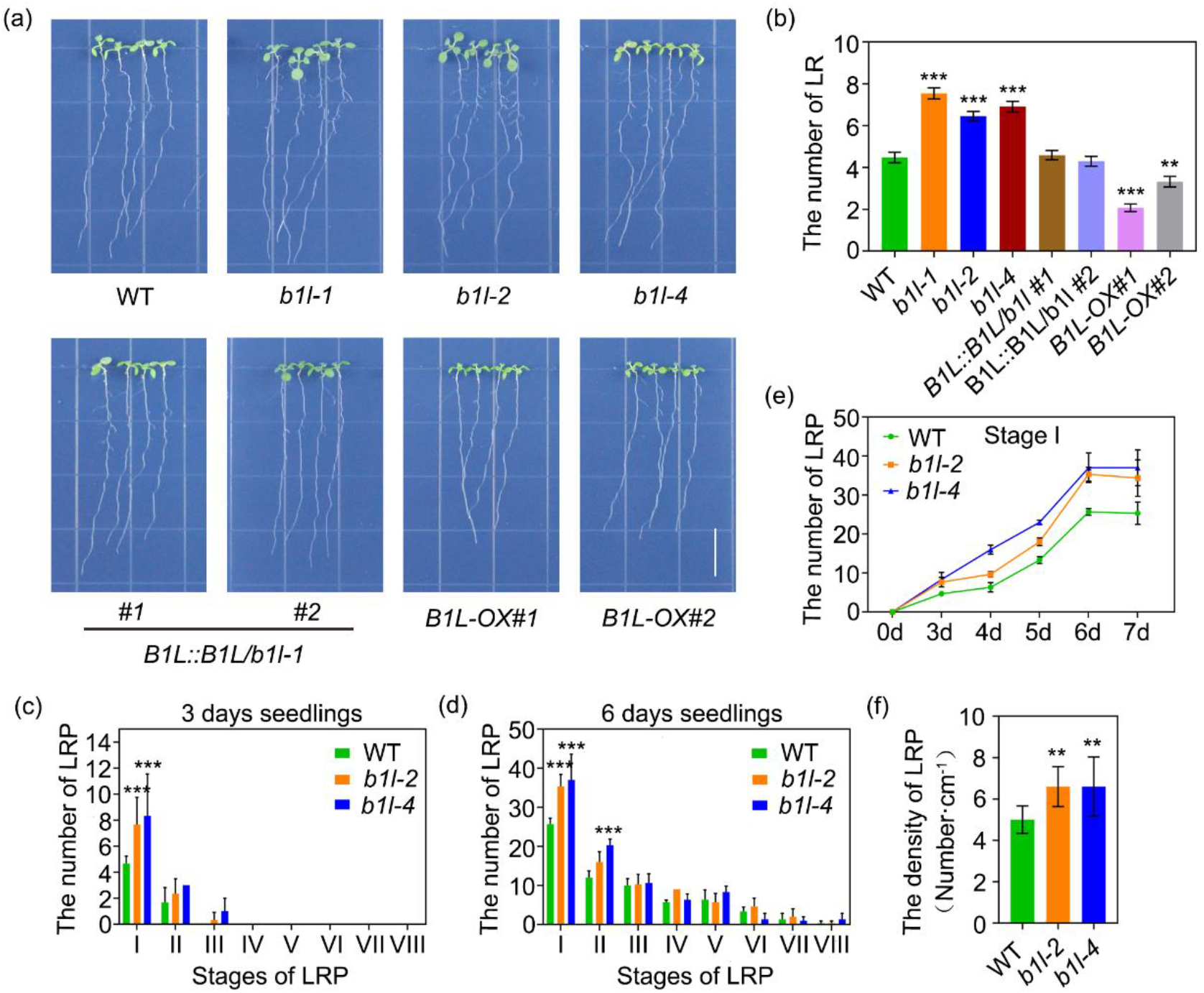
B1L regulates the lateral root initiation. (a) Root phenotypes of wild-type (WT), *b1l-1, b1l-2, b1l-4, B1L::B1L-3×Flag/b1l#1*, *B1L::B1L-3×Flag/b1l#2, B1L-OX#1* and *B1L-OX#2*. WT is Col-0.10 days old seedlings were used for photographs and measurements. Bars is 1.5cm. (b) The number of lateral roots in Figure 2A, seedlings. The values are means ± SD (n=44, *one-way ANOVA*, **, P < 0.01, ***, P < 0.001). (c, d) The distribution of lateral root primordia at I-VIII stages of WT, *b1l-2* and *b1l-4* seedlings at 3 days and 6 days, respectively. The values are means ± SD (n=3, *two-way ANOVA*, ***, P < 0.001 compared with WT at each sate). (e) The number of stage I lateral root primordia in WT, *b1l-2* and *b1l-4* seedlings at 0-7 days. The values are means ± SD (n=3, two-way ANOVA, **, P < 0.01, ***, P < 0.001 compared with WT at each time). (f) The lateral root primordia density of WT, *b1l-2* and *b1l-4* seedlings at 10 days. The values are means ± SD (n=10, one-way ANOVA, **, P < 0.01 compared with WT).

In order to reveal the mechanism underlying B1L-mediated regulation of lateral root development, we examined the differences in the eight developmental stages of lateral root primordia in the WT, *b1l-2*, and *b1l-4*. No growth defects were observed at any stage of lateral root primordia development in *b1l-2* and *b1l-4* (Fig. S3). However, our data suggested that the number of lateral root primordia at stage I and stage II in *b1l-2* and *b1l-4* were significantly higher than the corresponding number in the WT (Fig. 2c-e), and the density of lateral root primordia in *b1l-2* and *b1l-4* was clearly greater than that in the WT (Fig. 2f). These results indicated that B1L was involved in the initiation of lateral root primordia.

### B1L is involved in PIN-mediated auxin efflux

Lateral root primordia initiation is well known to depend on local high auxin levels; furthermore, auxin is known to play a crucial role in all stages of lateral root development (Stoeckle *et al*., 2018; Vilches-Barro and Maizel, 2015); therefore, we investigated whether B1L affects auxin signal during lateral root development. The auxin response reporter *DR5::GUS* was expressed in a similar pattern in WT and *b1l-1*. As the GUS staining showed, DR5::GUS was higher expressed at stage I in lateral root primordia of *b1l-1* than that in WT(Fig. 3). Moreover, IAA treatment at 0.05 and 0.1 μM significantly increased the number of lateral roots in WT and *b1l-1* plants (Fig. 4a, b). Interestingly, the significant difference in the lateral root phenotypes was eliminated between the WT and *b1l-1* treated with 0.05 and 0.1 μM IAA (Fig. 4a, b). These findings indicated that the lateral root phenotypes of the WT and *b1l-1* can be attributed to auxin levels.

**Figure 3.**
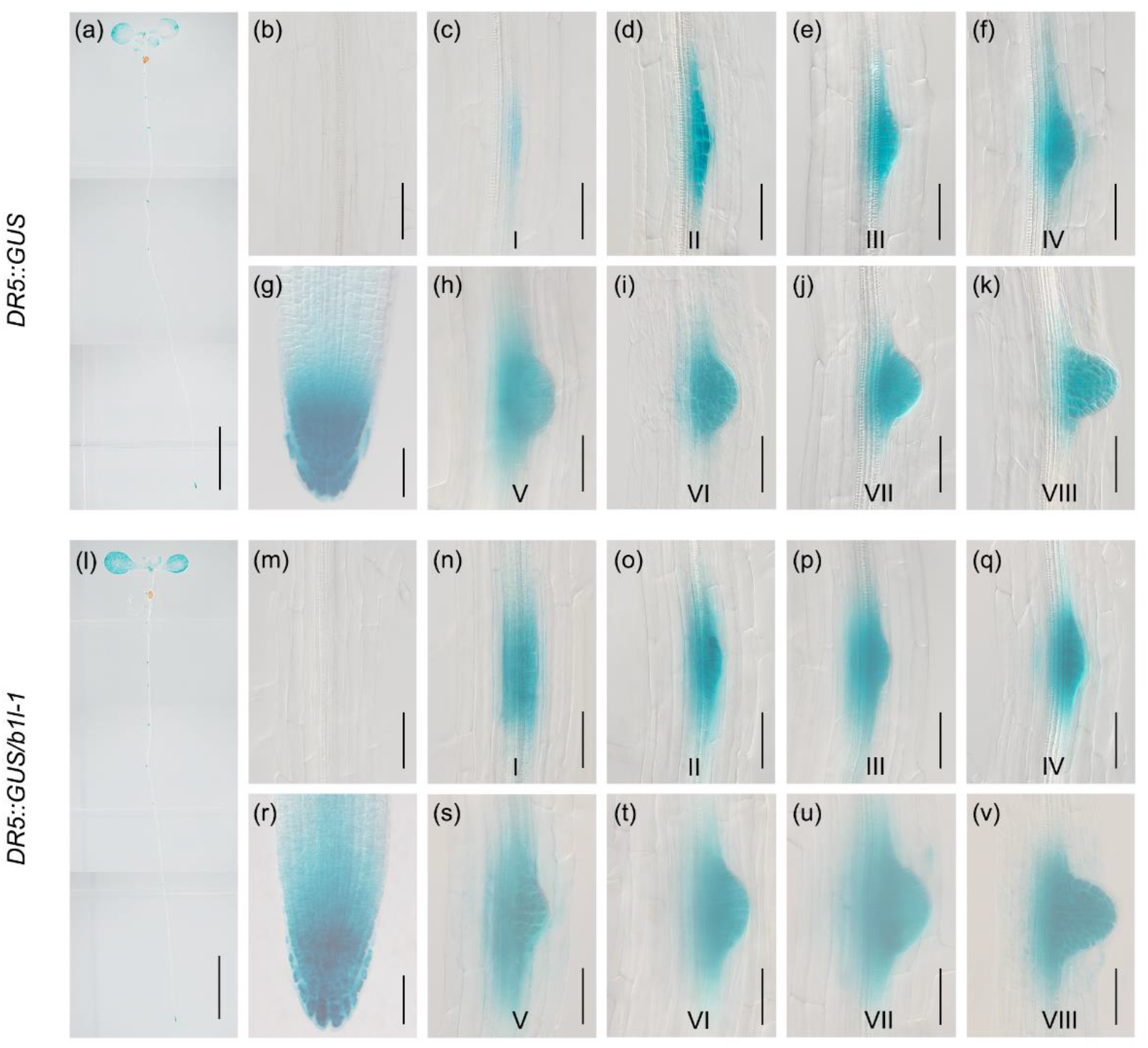
The auxin levels are increased in primary root and lateral root primordia of *b1l-1* mutant. The expression of *DR5::GUS* in whole plant (a, l), vascular cylinder (b, m), primary root (g, r) and eight different stages of lateral root primordium (c-k) and (n-v) of WT and *b1l-1* mutant, respectively. 10 days old seedlings were used for GUS staining. I-VIII are eight development stages of lateral root primordia. Images were obtained by DICM, bars in (a, l) are 5mm, and bars in (b-v) are 50μm.

**Figure 4.**
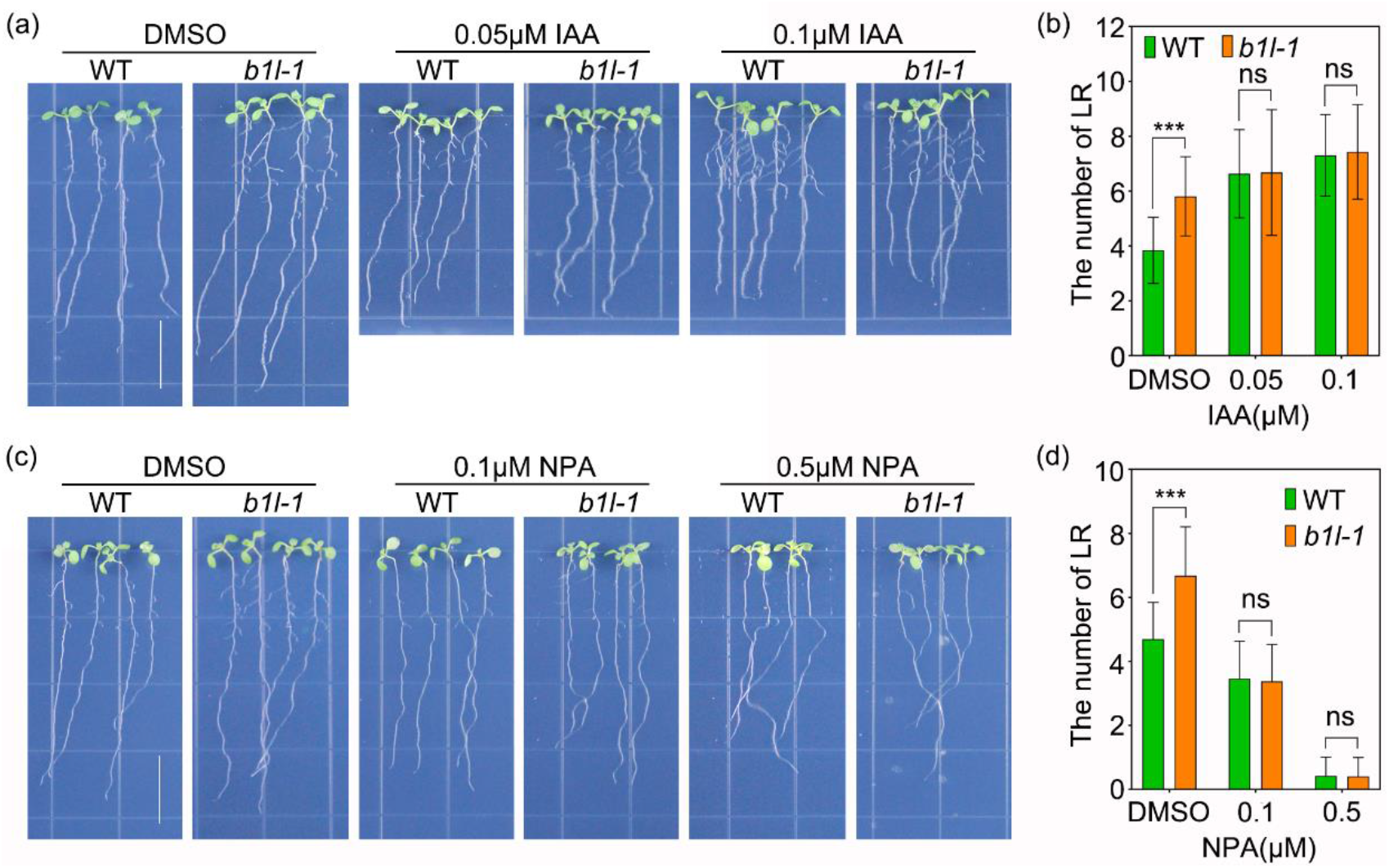
B1L regulates the auxin level by affecting the auxin efflux. (a, b) Phenotypic analysis of lateral root in WT and *b1l-1* seedlings treated with IAA. The values are means ± SD (n≥58, Student’s *t-test*, ***, P < 0.001 compared with WT). (c, d) Phenotypic analysis of lateral root in WT and *b1l-1* seedlings treated with NPA. Bar is 1.5 cm. The values are means ± SD (n≥83, Student’s *t-test*, ***, P < 0.001 compared with WT).

As the level of auxin in plants is mainly determined by its biosynthesis and transport (Du and Scheres, 2018), first, we investigated whether B1L is involved in auxin biosynthesis. Treatment with auxin biosynthesis inhibitor L-kynurenine (L-kyn) at concentrations of 0.1 and 0.5 μM significantly inhibited the number of lateral roots in both the WT and *b1l-1* plants, however, the difference in lateral root phenotype between WT and *b1l-1* was still remained (Fig. S4 a, b). Furthermore, auxin biosynthesis genes were not upregulated in the *b1l-1* mutant at the transcriptional level (Fig. S5a). Therefore, we concluded that B1L may be not involved in auxin biosynthesis.

Second, to assess whether B1L affects auxin transport, auxin influx inhibitor 2-naphthoxyacetic acid (2-NOA) and auxin efflux inhibitor N-naphthylphthalamic acid (NPA) were used. Both 2-NOA and NPA significantly reduced the number of lateral roots in both WT and *b1l-1* plants (Fig. S4c, d; Fig. 4c, d). Moreover, the lateral root number still differed between 2-NOA-treated WT and *b1l-1* plants; however, this difference was eliminated upon treatment with NPA. This result indicated that B1L affects the auxin efflux. To further determine the impact of B1L on auxin efflux, we introduced *AUX1::AUX1-YFP, PIN1::PIN1-GFP*, and *PIN3::PIN3-GFP into b1l-1* to observe the expression patterns of the influx carrier AUX1 and efflux carrier PIN family in the roots of WT and *b1l-1* plants. As expected, we observed that the levels of PIN1-GFP increased in *b1l-1* lateral root primordia at all eight stages (Fig. 5a), and the levels of PIN3-GFP increased in the lateral root primordia of *b1l-1* from stage I to stage III (Fig. 5b). In addition, no pronounced changes were observed in AUX1-YFP levels between WT and *b1l-1* (Fig. S6). Furthermore, at the transcriptional level, PINs did not increase in the *b1l-1* mutant (Fig. S5b). Taken together, these findings suggest that B1L regulates polar auxin transport by its involvement in PIN-mediated auxin efflux.

**Figure 5.**
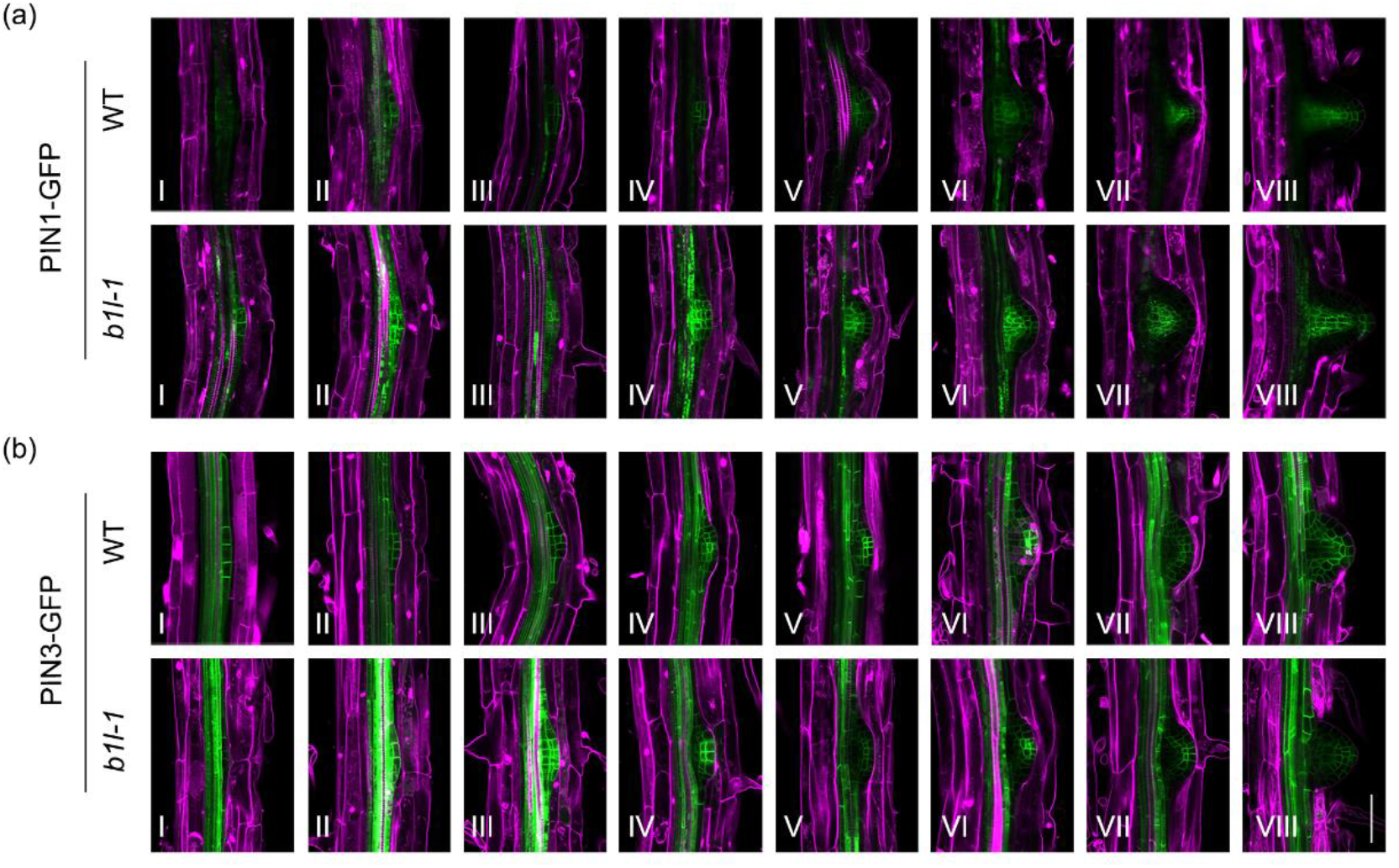
The auxin efflux carrier PIN1-GFP and PIN3-GFP are increased in lateral root primordia of *b1l-1* mutant. (a) The expression of *PIN1::PIN1-GFP* in eight development stages of lateral root primordia of WT and *b1l-1* mutant. (b) The expression of *PIN3::PIN3-GFP* in eight development stages of lateral root primordia of WT and *b1l-1* mutant. I-VIII are eight development stages of lateral root primordia. Images were obtained by confocal microscope, bar is 50μm.

### B1L interacts with the exocyst complex

To explore the mechanism whereby, B1L affects PIN levels, we identified potential proteins that interact with B1L using co-immunoprecipitation (co-IP) and liquid chromatography-tandem mass spectrometry (LC-MS/MS). Anti-Flag antibodies conjugated to agarose beads were used to immunoprecipitate B1L-3×Flag and its interacting proteins from root proteins of *B1L::B1L-3×Flag* and WT, which grew 7 d on 1/2 MS medium plates. Five bands, with the exception of heavy and light chains, were observed by SDS-PAGE (Fig. S7a) and then analyzed by LC-MS/MS. Among these, 888 proteins that were larger than or equal to the two unique peptides, were identified. Next, GO enrichment analysis revealed that these proteins were highly enriched in cellular processes (43.89%), metabolic processes (33.95%), biological regulation (8.27%), localization (7.21%), and response to stimulus (5.45%) (Fig. S7b). Interestingly, 37 vesicle-mediated transport proteins and 11 exocyst subunits were observed in the cellular process (Fig. S7c; Fig. 6a). In addition, EXO70B1 was previously identified in our yeast two-hybrid (Y2H) screening assay using B1L-pGBKT7 as bait. These results indicated that B1L interacts with the exocyst complex.

**Figure 6.**
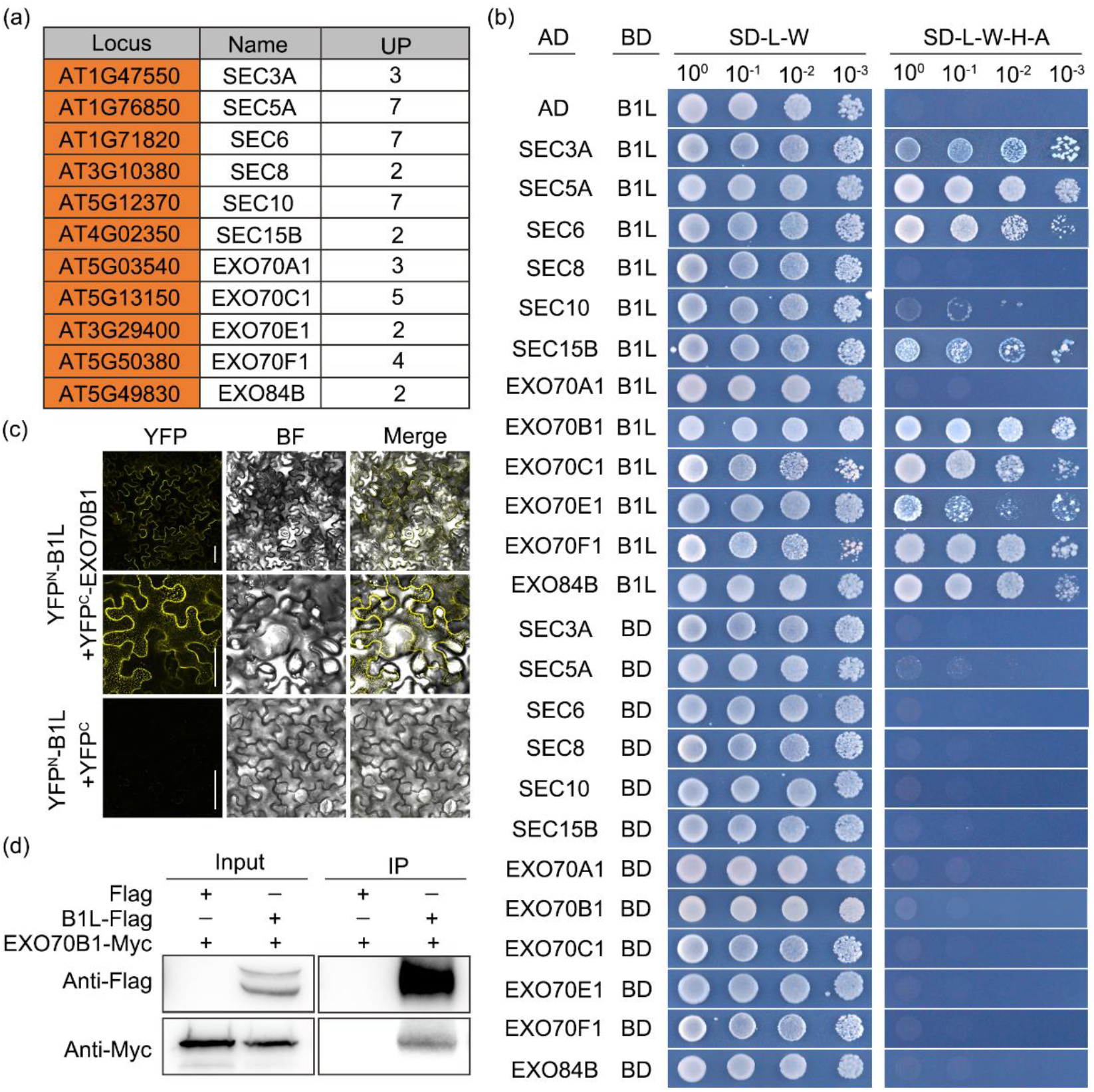
B1L interacts with exocyst complex. (a) 11 exocyst subunits were identified by LC-MS/MS in eluants immunoprecipitated by anti-Flag agarose beads. UPs are unique peptides. (b) The interactions among B1L and exocyst subunits were verified by yeast two-hybrid (YH2) assays. The empty pGADT7 and pGBKT7 vectors were used as negative controls. (c) BiFC analysis of the interaction between B1L and EXO70B1 in *N. benthamiana*. Bars are 50μm. (d) Co-IP confirm the interaction between B1L and EXO70B1. *35s::EXO70B1-Myc* were co-transfected with *35s::B1L-Flag* or *Flag* empty vector in *N. benthamiana* leaves. Total proteins were immunoprecipitated with anti-Flag agarose beads. B1L-Flag was detected with anti-Flag antibody and immunoprecipitated EXO70B1-Myc was detected with anti-Myc antibody.

To confirm the potential interaction between B1L and the exocyst, we first examined the physical interactions of B1L with the exocyst subunits using the Y2H assay. We found that B1L interacted with SEC3A, SEC5A, SEC6, SEC10, SEC15B, EXO70B1, EXO70C1, EXO70E1, EXO70F1, and EXO84B in yeast (Fig. 6b). Subsequently, we performed a bimolecular fluorescence complementation (BiFC) assay. Transient co-expression of the fusion genes *YFP^N^-B1L* and *YFP^C^-EXO70B1* in *N. benthamiana* leaves resulted in YFP fluorescence (Fig. 6c). Interestingly, YFP fluorescence was localized not only in the plasma membrane, but in vesicle-like structures as well (Fig. 6c). Co-immunoprecipitation (Co-IP) was used to confirm this interaction. Constructs *35S::B1L-Flag* and *35S::EXO70B1-Myc* were transiently co-expressed in *N. benthamiana* leaves. Total leaf protein extracts were immunoprecipitated with anti-Flag agarose beads, and the precipitated proteins were detected using anti-Myc antibodies. As expected, a band of the size of EXO70B1-Myc was detected in the anti-Flag agarose precipitated fractions (Fig. 6d). These results strongly indicated a direct interaction between B1L and the exocyst *in vivo*.

### B1L is involved in the recycling of PIN2

The plant exocyst complex facilitates tethering between transport vesicles and the plasma membrane, and plays an important role in auxin efflux carrier PINs recycling and polar auxin transport (Tan *et al*., 2016; Drdová *et al*., 2013; Žárský *et al*., 2013). Thus, we investigated whether the accumulation of PINs in *b1l-1* was due to a defect in recycling and trafficking. To this purpose, we examined PIN2-GFP trafficking in the WT and *b1l-1* roots by treating them with Brefeldin A (BFA) and cycloheximide (CHX). BFA is a reversible vesicle trafficking inhibitor that causes the aggregation of PIN2 and formation of BFA bodies (Mayers *et al*., 2017). CHX is an inhibitor of eukaryotic translation that can eliminate the differences in PIN2 synthesis (Schneider-Poetsch *et al*., 2010). BFA bodies accumulated both in the WT and *b1l-1* mutant of root epidermal cells treated with BFA+CHX after 60 min (Fig. 7a, b). The number of BFA bodies in the WT markedly decreased compared to that in *b1l-1* after BFA washout at 60 and 90 min. These findings confirmed our hypothesis that *b1l-1* exhibits defects in PIN2 exocytosis. Thus, we demonstrated that B1L is involved in the recycling of PIN2.

**Figure 7.**
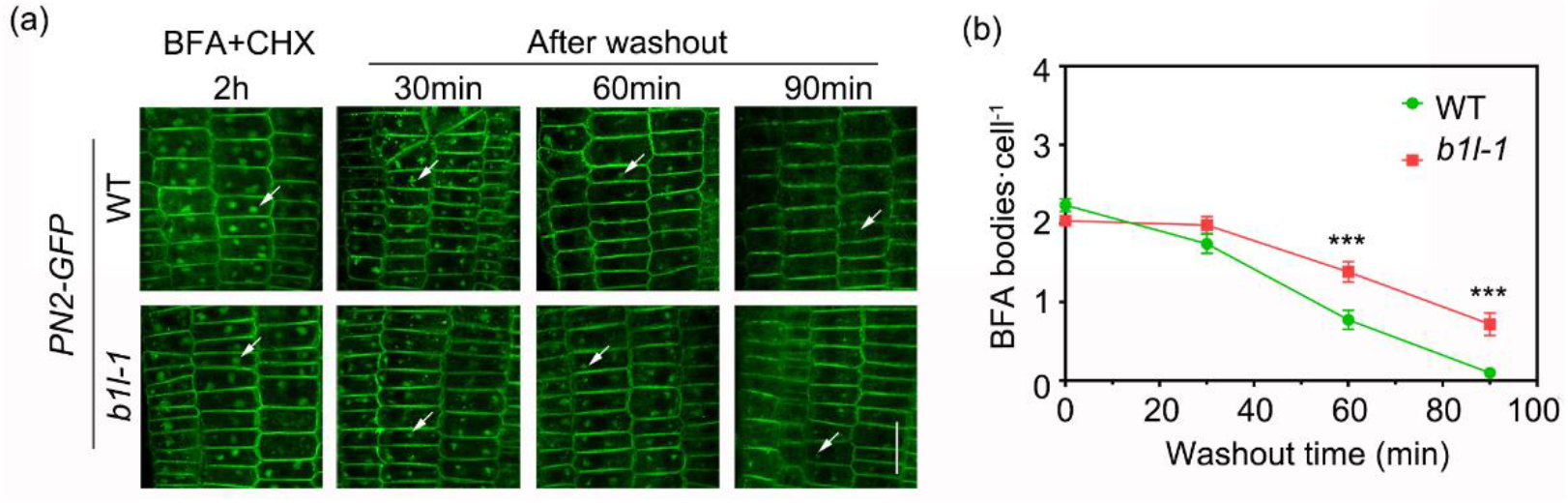
B1L is involved in the recycling of PIN2. (a) *b1l-1* mutants showed defect in PIN2-GFP exocytosis. *PIN2-GFP* and *PIN2-GFP/b1l-1* seedlings were treated with 50μM CHX for 60min and then were treated with 50μM BFA and 50mM CHX for 2h. The BFA washout with 1/2 M and imaged at 0, 30, 60, and 90min by confocal microscope. White arrowheads are PIN2-GFP-labeled BFA bodies. Bar is 20μm. (b) The number of PIN2-GFP-labeled BFA bodies per cell after BFA washout at indicate times. The values are means ± SEM (n≥29, *two-way ANOVA*, ***, P < 0.001 compared with WT at each time).

### *b1l-1/exo70b1-1* double mutant shows a significantly increased lateral root formation phenotype

At last, we investigated the genetic interactions between *B1L* and *EXO70B1*. The *b1l-1/exo70b1-1* double mutant was generated by crossing *b1l-1* and *exo70b1*. The lateral root phenotype of *exo70b1* was similar to that of *b1l-1*, which exhibited a greater number of lateral roots than the WT (Fig. 8). Interestingly, the lateral root phenotype of *b1l-1* and *exo70b1* significantly increased in the *b1l-1/exo70b1-1* double mutant (Fig. 8). These results suggested that B1L interacts with EXO70B1 to regulate lateral root formation, cooperatively.

**Figure 8.**
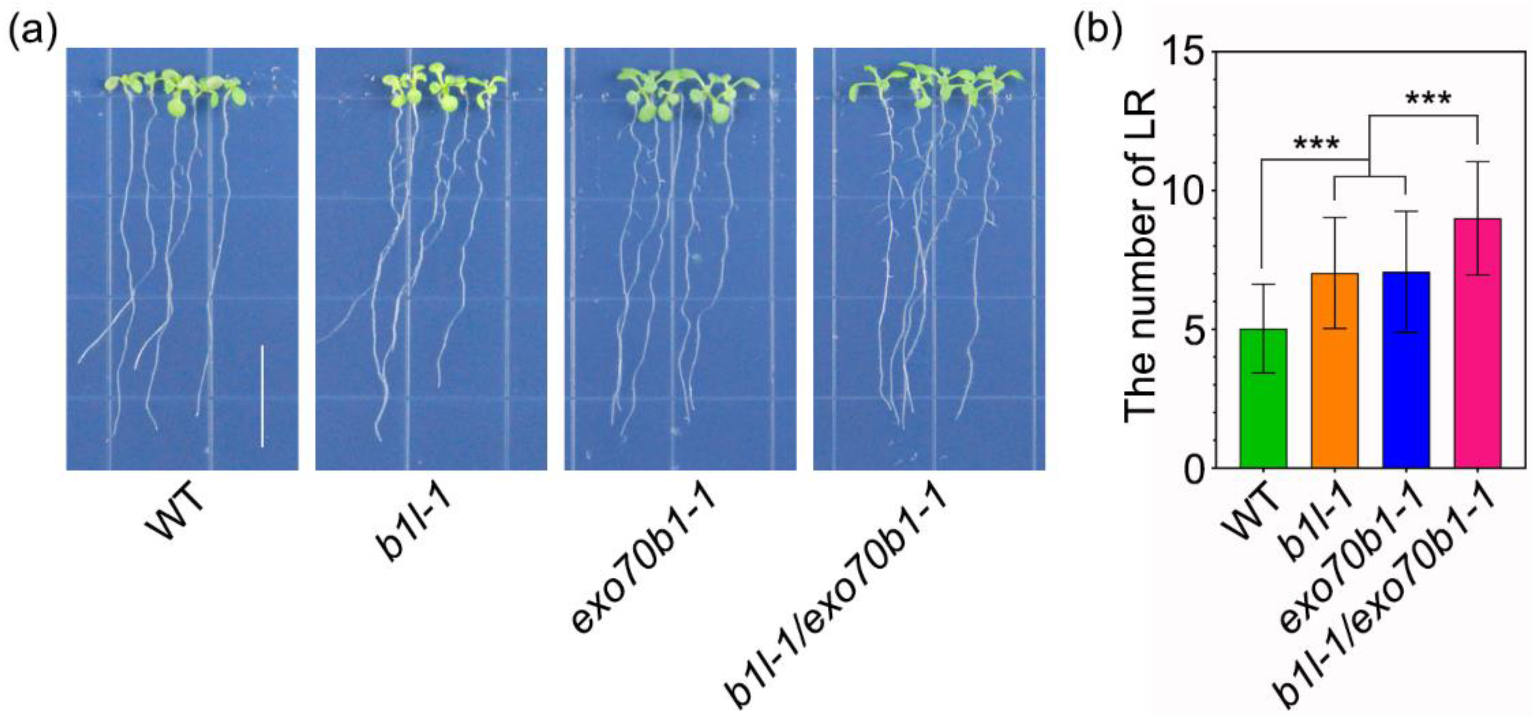
B1L cooperated with EXO70B1 to regulate the lateral roots development. (a) Lateral root phenotypes of WT, *b1l-1, exo70b1-1* and *b1l-1/exo70b1-1*. Bar is 1.5 cm. (b) The number of lateral roots in Fig.8a. The values are means ± SD (n≥60, *one-way ANOVA*, ***, P < 0.001 compared with WT or single mutants).

## DISCUSSION

Our previous studies showed that B1L is involved in the regulation of freezing tolerance in plants (Chen *et al*., 2019; Chen *et al*., 2020); however, other biological functions of B1L remain unclear. In this study, we revealed a novel biological function and the elegant underlying molecular mechanism of B1L-mediated regulation of lateral root formation. The expression pattern of B1L plays an important role in the development of primary and lateral roots (Fig. 1). Genetic and cellular analyses showed that B1L regulates lateral root development and lateral root primordia initiation (Fig. 2 and Fig. S3). In fact, the presence of more lateral roots is not always advantageous for plants. Lateral roots develop only at specific tissue positions and time points to accomplish root development for water and nutrient uptake, because every increase in the number of organs requires additional energy(Péret *et al*., 2009; Möller *et al*., 2017). Therefore, we consider that the larger number of lateral roots due to the absence of B1L is not necessarily beneficial to healthy plant development and growth.

Lateral root primordia initiation occurs in xylem pole pericycle cells located in the basal meristem or root elongation zone (Motte *et al*., 2019; Péret *et al*., 2009). The pericycle cells are stimulated by a local auxin threshold level and become lateral root-founder cells that subsequently undergo anticlinal and asymmetric divisions to generate a single-layer primordium referred to as stage I(Lavenus *et al*., 2013; Vilches-Barro and Maizel, 2015; Péret *et al*., 2009; Motte *et al*., 2019). Therefore, auxin is widely believed to play a crucial role in regulating lateral root primordia initiation (Du and Scheres, 2018; Lavenus *et al*., 2013; Bellini *et al*., 2014). In the present study, higher auxin levels were found in the lateral root primordia of the *b1l* mutant (Fig. 3), and exogenous auxin treatment confirmed that lateral root phenotype correlated closely with auxin level (Fig. 4a, b). The local auxin threshold is maintained by auxin influx and efflux carrier-mediated (i.e., AUX1 and PINs) transport. Our data showed that the differences in lateral root phenotype between WT and mutant roots were eliminated upon treatment with the auxin efflux inhibitor NPA (Fig. 4c, d). Furthermore, the *b1l* mutant exhibited higher levels of PINs (Fig. 5). This finding indicated that B1L regulates auxin efflux mainly by influencing the level of PINs.

Co-IP/LC-MS/MS analysis is a powerful tool to unravel the potential roles of B1L in regulating the level of PINs. We identified 12 exocyst complex subunits that potentially interact with B1L (Fig. 6a; Fig. S7). These interactions were confirmed by performing Y2H, BiFC, and Co-IP assays (Fig. 6b-d). Our findings indicated that B1L interacts with the tethering exocyst complex composed of eight subunits (SEC3, SEC5, SEC6, SEC8, SEC10, SEC15, EXO70, and EXO84), which facilitates vesicle contact with the target membrane(Žárský *et al*., 2013; Mei and Guo, 2018; Wu and Guo, 2015). As the previous studies have shown, different subunits of exocyst complex are involved in different biological processes. SEC3A, SEC5, SEC6, SEC8 and SEC15A are reported to be involved in pollen development(Bloch *et al*., 2016; Hála *et al*., 2008). Their mutants showed defects in pollen germination and pollen tube growth. The absence of EXO84B resulted in defective cytokinesis and a large accumulation of vesicles in the cytoplasm, suggesting that exocyst complex mediated vesicle transport is involved in cell plate formation(Fendrych *et al*., 2010). Exocyst mutants *exo70a1* and *sec8* showed defects in the recycling of PIN1 and PIN2 proteins(Drdová *et al*., 2013). In addition, a recent study revealed that the *sec6* mutant exhibited the same phenotype as *exo70a1* and *sec8* mutants in PIN1 and PIN2 protein recycling (Tan *et al*., 2016). Thus, the exocyst is widely considered to be involved in regulating PINs and recycling other plasma membrane proteins (Gao *et al*., 2008; Zhou and Luo, 2018; Naramoto, 2017). PIN2 is located in epidermal cells in roots and is extensively used in BFA treatment assays to investigate vesicle trafficking (Zhou and Luo, 2018; Pavel Křeček *et al*., 2009; Mayers *et al*., 2017). In this study, the *b1l* mutant exhibited a defect in PIN2-GFP recycling (Fig. 7a, b), which indicated that B1L functions in exocytic vesicle trafficking of PINs through interaction with the exocyst. Although we investigated PIN2-GFP recycling, instead of PIN1-GFP and PIN3-GFP, this did not affect our conclusions.

Another interesting finding in this study was that EXO70B1 was reported to be involved in lateral root formation. As the previous studies have shown, EXO70B1 is essential for autophagosome formation and immune response in Arabidopsis(Zhao *et al*., 2015; Kulich *et al*., 2013). Recent studies revealed that EXO70B1 and EXO70B2 regulate FLS2 homeostasis at the plasma membrane(Wang *et al*., 2020). However, there is no study to report that whether EXO70B1 is involved in lateral root development. In this study, we found *exo70b1* mutant showed a greater number of lateral roots phenotype compared with WT (Fig. 8a, b). Interestingly, the double mutant *b1l-1/exo70b1-1* enhanced the lateral roots phenotype. These results suggest that B1L and EXO70B1 may play functionally redundant roles and cooperatively regulate lateral roots development.

In summary, we demonstrated that B1L is involved in regulating auxin levels by interacting with the exocyst to function in the exocytic vesicle trafficking-mediated PIN recycling in WT root cells (Fig. S8). The absence of B1L in *b1l* mutant seemingly results in abnormal PIN recycling that leads to a higher local auxin level, which in turn facilitates the initiation of lateral root primordia. This study improves our understanding of the highly sophisticated processes involved in exocytic vesicular trafficking-mediated polar auxin transport and lateral root initiation in plants.

## EXPERIMENTAL PROCEDURES

### Plant materials and growth conditions

Wild-type (WT), mutants and transgenic plants used in this study are all Col-0 background of *Arabidopsis thaliana*. The T-DNA insertion mutants *b1l-1* (SALK_020993) and *b1l-2* (SALK_019913) were obtained from the Arabidopsis Biological Resource Center. The T-DNA insert information are showed in Fig. S1a. *b1l-4* mutant is the transgene-free gene editing mutant obtained by using the CRISPR/Cas9 system. *b1l-4* absents 96bp from 13bp to 108bp after initiator codon ATG. The CRISPR/Cas9 system vectors were kindly provided by the Qi-Jun Chen laboratory of China Agricultural University, and the protocols performed as previously described(Wang *et al*., 2015). The primer sequences that were used for CRISPR/Cas9 system are listed in Table S1. The primer sequences that were used for identification of *b1l-1, b1l-2 and b1l-4* are listed in Table S2 and the results are showed in Fig.S1. Transgenic Arabidopsis plants expressing GUS driven by the *B1L* promoter (*B1L::GUS*), B1L-3×Flag driven by the *B1L* promoter (*B1L::B1L-3×Flag/b1l*) and YFP-B1L driven by the CaMV35S promoter (*B1L-OX*) were previously described(Chen *et al*., 2019).

Transgenic Arabidopsis plants expressing GUS driven by the *DR5* promoter (*DR5::GUS*), PIN1-GFP driven by the *PIN1* promoter (*PIN1::PIN1-GFP*), PIN2-GFP driven by the *PIN2* promoter (*PIN2::PIN2-GFP*), PIN3-GFP driven by the *PIN3* promoter (*PIN3::PIN3-GFP*), AUX1-YFP driven by the *AUX1* promoter (*AUX1::AUX1-YFP*) were kindly provided by the Guang-Qin Guo laboratory. The above plants were crossed by *b1l-1* mutant and generated *DR5::GUS/ b1l-1, PIN1::PIN1-GFP/b1l-1, PIN2::PIN2-GFP/b1l-1*, *PIN3::PIN3-GFP/b1l-1* and *AUX1::AUX1-YF/b1l-1*, respectively.

All plants were grown on 1/2 Murashige and Skoog (MS) medium plates in an artificial climate chamber (RXZ-500; Ningbo Jiangnan, China) with 16 h light and 8 h dark at 22°C.

### Chemical treatment assays

The seedlings grown 3 days after germination on MS medium plates were transferred on the 1/2 MS medium plates contained different concentrations of medicines. The phenotypes were analyzed and photographed after 7 days. The medicines included the auxin Indole-3-acetic acid (IAA, 0.05 and 0.1 μM), the auxin synthesis inhibitor L-kynurenine (L-kyn, 0.1 and 0.5 μM), auxin influx inhibitor 2-naphthoxyacetic acid (2-NOA, 0.02 and 0.05 μM) and auxin efflux inhibitor N-Naphthylphtalamic acid (NPA, 0.1 and 0.5 μM)

### Lateral root primordia analysis

The observation of lateral root primordia was performed as previously described(Xun *et al*., 2020). The roots of 10 days old seedlings were dipped with 0.4 M HCl in 20% methanol for 15min at 57°C, then transferred into 7% NaOH and 7% hydroxylamine-HCl in 60% ethanol for 15min at room temperature. The roots were washed 5 min with 40%, 20% and 10% ethanol, respectively, and then dipped in 5% ethanol and 25% glycerol solution for 15min. At last, the roots were stored in 50% glycerol solution. The roots and lateral root primordia were analyzed and photographed by a research grade automatic positive fluorescence microscope (Axio Imager.Z2; Zeiss, Germany).

### GUS staining

The GUS staining was performed as previously described(Xun *et al*., 2020). 10 days old seedlings of *B1L::GUS, DR5::GUS* and *DR5::GUS/ b1l-1* were incubated in GUS staining solution at 37°C for 6 h, then the whole plants were rinsed with different concentrations of alcohol, and the lateral root primordia were cleared with hydroxylamine-HCl as described above. A Stereo Fluorescence Microscope (Discover.v20; Zeiss, Germany) was used to analyze and photograph for whole plants. The roots and lateral root primordia were analyzed and photographed by a research grade automatic positive fluorescence microscope (Axio Imager.Z2; Zeiss, Germany).

### Image analysis with LSCM

For protein localization of PIN1-GFP, PIN3-GFP and AUX1-YFP experiments, the roots of 10 days old seedlings were stained with 10mg·ml^-1^ propidium iodide (PI) for 6 min. A laser-scanning confocal microscopy (LSCM) (LSM880; Zeiss, Germany) was used to analyze and photograph. GFP or YFP was excited with the 488nm laser, and PI was excited with the 561nm laser. The emission of GFP or YFP and PI was detected between 500-530 nm and 570–670 nm by a multichannel detector with filters, respectively.

For protein localization of SEC12-YFP and SYP32-YFP experiments, the roots of 10 days old seedlings were directly analyzed and photographed by LSCM (LSM880; Zeiss, Germany) at a single channel detector.

For PIN2-GFP trafficking experiments, seedlings were firstly treated with 50 μM CHX (Sigma-Aldrich) in 1/2 MS liquid medium for 1 hour. Then the seedlings were transferred into 1/2 MS liquid medium contained 50 μM CHX and 50 μM BFA (Sigma-Aldrich) for 2 hours. After treatment, the seedlings were washout with MS liquid medium for 0, 30, 60 and 90 minutes. Seedlings were imaged by CLSM (LSM880; Zeiss, Germany) at indicated times.

### Total RNA extract and qRT-PCR assay

The roots of 7 days old seedlings were collected to extract total RNA using RNA prep pure plant kit (TIANGEN). The first strand cDNA was synthesized from 1 μg of total RNA using the Hifair^®^ III 1st Strand cDNA Synthesis SuperMix kit (YEASEN, 11141ES10) according to the manufacturer’s instructions.

For qRT-PCR, it was performed by ABI Real-Time PCR Detection System (ABI, Q5) using the Hieff^®^ qPCR SYBR Green Master Mix (Low Rox Plus) (YEASEN, 11202ES08). Three biological replicates and four technical replicates were performed for each sample. *AtUBC21* (At5g25760) was used as the reference control(Cai *et al*., 2017). The primer sequences used for qRT-PCR are listed in Supplemental Table S3.

### Yeast two-hybrid assay

Yeast two-hybrid assays were performed a s previously described(Chen *et al*., 2019). The CDS of B1L was cloned into the pGBKT7-GW and the CDSs of exocyst subunits were cloned into the pGADT7-GW vector using gateway cloning system, respectively. The primer sequences are listed in Table S4. The B1L-pGBKT7 vector and pGADT7 fused exocyst subunit vector were co-transformed into AH109 strain. The interactions between B1L and exocyst subunits were determined by observing the growth status of positive clones on media lacking leucine (Leu), tryptophan (Trp), histidine (His), and adenine (Ade).

### BiFC assay

Bimolecular fluorescence complementation (BiFC) assay was performed as previously described(Chen *et al*., 2019). The CDSs of B1L and EXO70B1 were cloned into the PNYFP-X and PCCFP-X vectors using gateway cloning system, respectively. The primer sequences are listed in Table S4. Transient co-expressed the fusion genes *YFP^N^-B1L* and *YFP^C^-EXO70B1* in *N. benthamiana* leaves. The interaction was analyzed and photographed by LSCM (Leica; SP8, Germany)

### IP/ Co-IP assay

The IP assay was performed as previously described(Chen *et al*., 2019). In brief, total protein was extracted from roots of 7 days old WT and *B1L::B1L-3×Flag* seedlings using IP buffer (50Mm Tris-HCl, pH 7.6, 150mM NaCl, 10% Glycerol, 0.1% NP-40 and 1×Cocktail). Protein extracts were added into 50μl agarose beads conjugated anti-Flag antibody (M20018L; Abmart, China), and the mix was incubated with gentle shaking for 6 h at 4 °C. The beads were washed with IP buffer five times, then added 50μl 1 ×loading buffer to boil 10 minutes in water. The beads were centrifuged 1 min at 12000rpm and the supernatant was used for running SDS-PAGE.

For Co-IP assay, the CDSs of B1L and EXO70B1 were cloned into the 35S-GATWAY-3×Flag vector and the 35S-GATWAY-3×Myc vector using gateway cloning system, respectively. The primer sequences are listed in Table S4. The *35S::B1L-Flag* and *35S::EXO70B1-Myc* constructs were transiently co-expressed in *N. benthamiana* leaves. *35S::Flag* and *35S::EXO70B1-Myc* constructs were transiently co-expressed as control. Total protein was extracted from leaves using IP buffer, and the IP was performed as described above. Anti-Flag antibody (M20008M; Abmart, China) and anti-Myc antibody (ab32072; Abcam, UK) were used to detect B1L-Flag and EXO70B1-Myc, respectively.

### Western blot

Proteins were extracted as described above and separated on SDS-PAGE (12%), and then transferred to a PVDF membrane (0.22μm; Millipore, America) at 200 mA for 1.5h in transfer buffer. The PVDF membrane was blocked in 5% Non-Fat Powdered Milk (A600669; Sangon Biotech, China) 1h at 22°C. After incubation with primary antibodies and secondary antibodies respectively, the PVDF membrane was treated with enhanced chemiluminescent reagent (NCM Biotech, P10200, China) and then imaged using chemiluminescence imaging analysis system (BG-gdsAUTO 710 MINI; Baygene biotech, China).

### LC-MS/MS

LC-MS/MS analysis was performed as previously described which mainly included alkylation, tryptic digestion, mass spectrometric and database searching(Wang *et al*., 2013; Zhang *et al*., 2020). The gels of protein bands were cut into cubic pieces and washed with 50% acetonitrile in 25 mM NH_4_HCO_3_ until the Coomassie brilliant blue was disappeared. The gels were dehydrated by acetonitrile and vacuum dried. Next, the gels were alkylated with 10mM TCEP and 40Mm CAA, and dehydrated by acetonitrile again. The gels were digested with 0.01mg·ml^-1^ trypsin in 100Mm NH_4_HCO_3_ solution 8-24h at 37°C. The gels were putted into new tubes and extracted with 0.1% TFA in 50% acetonitrile. The extracts were mixed with digest solution and vacuum dried. 0.1% FA was added to dissolve polypeptides. The solutions were centrifuged at 14000rpm for 30min. The supernatants were analyzed by liquid chromatography electrospray tandem mass spectrometry (Orbitrap Fusion Lumos Easy 1200 Nano LC, Thermofisher). The results were analyzed using thermo proteome discoverer 2.1.1.21 software to search *Arabidopsis* protein database.

## ACCESSION NUMBERS

Sequence data from this article can be found in The Arabidopsis Information Resource (http://www.arabidopsis.org/). Accession number: *B1L*(AT1G18740).

## Acknowledgements

We thank Guang-Qin Guo (Lanzhou University, Lanzhou, Gansu, China) for providing the *DR5::GUS, PIN1::PIN1-GFP, PIN3::PIN3-GFP* and *AUX1::AUX1-YFP* transgenic plants, Qi-Jun Chen (China Agricultural University, Beijing, China) for providing the CRISPR/Cas9 system vectors, and the Core Facility for Life Science Research (Lanzhou University) for technical assistance. This work was supported by the Key Program National Natural Science Foundation of China (41830321), the National Natural Science Foundation of China (31770432, 32071482) and the Fundamental Research Funds for the Central Universities (lzujbky-2020-31).

## AUTHOR CONTRIBUTIONS

HZ, X-LY and L-ZA conceived the project. B-XC, GY, J-HC, TC, RS, C-CL and JJ performed the experiments. GY analyzed the data and wrote the manuscript. HZ, X-LY and L-ZA examined the data and manuscript.

## SUPPORTING INFORMATION

Additional Supporting Information may be found in the online version of this article.

**Figure S1** The information of *b1l* mutants.

**Figure S2** The length of primary roots are no obvious difference among WT, *b1l-1, b1l-2* and *b1l-4*.

**Figure S3** Morphology of lateral root initiation at different stages in WT, *b1l-2* and *b1l-4* mutants seedling roots.

**Figure S4** B1L is not involved in the auxin biosynthesis and the auxin influx.

**Figure S5** The auxin biosynthesis genes and *PINs* were not increased at the transcriptional levels in *b1l-1* mutant.

**Figure S6** The expression of *AUX1::AUX1-YFP* in eight development stages of lateral root primordia of WT and *b1l-1* mutant.

**Figure S7** The immunoprecipitation and LC-MS/MS analysis of proteins that potentially interact with B1L.

**Figure S8** The model for B1L in regulating lateral root development in Arabidopsis.

**Table S1** The primers of vectors construction for CRISPR/Cas9.

**Table S2** The primers for identification of *b1l-1, b1l-2* and *b1l-4* mutants.

**Table S3** The primers used for qRT-PCR in this study.

**Table S4** The primers of vectors construction for Y2H, BiFC and Co-IP assays.

